# clusterSeq: methods for identifying co-expression in high-throughput sequencing data

**DOI:** 10.1101/188581

**Authors:** Thomas J. Hardcastle, Irene Papatheodorou

**Affiliations:** Department of Plant Sciences, University of Cambridge, Downing Street, Cambridge CB2 3EA, United Kingdom; European Bioinformatics Institute (EMBL-EBI), Wellcome Genome Campus, Hinxton, Cambridge, CB10 1SD, UK

## Abstract

****Summary:**:** Identifying gene co-expression is a significant step in understanding functional relationships between genes. Existing methods primarily depend on analyses of correlation between pairs of genes; however, this neglects structural elements between experimental conditions. We present a novel approach to identifying clusters of co-expressed genes that incorporates these structures.

****Availability:**:** The methods are released on Bioconductor as the clusterSeq package (https://bioconductor.org/packages/release/bioc/html/clusterSeq.html).

## INTRODUCTION

Identifying patterns of gene expression across different conditions within an experiment is an important step in understanding possible co-expression or co-regulation and delineating complex relationships between genes. Typically, correlation methods are used to address this problem. Available methods, such as WGCNA (Langfelder and Horvath, 2008), employ correlation measures of gene expression across conditions to build connections across genes into a network and then identify network modules using clustering. However, structures within the data that derive from equivalent levels of expression between samples can confound measures of correlation.

Figure 1 illustrates the difficulties with correlation analyses that neglect these structures. Here genes are simulated from a log-normal distribution with standard deviation of 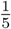, representing combined biological and technical variation. Simulations generate data for each gene over four replicates of four different experimental conditions A, B, C and D. Panel (a) shows the (log_2_)-expression of genes 1 & 2. For these genes, the expression of the genes clearly diverges over conditions A, B & C; however, the large magnitude change in expression in condition D gives a high Pearson correlation between the genes despite this divergence, implying a false positive identification of co-expression based on a Pearson correlation.

**Fig. 1.**
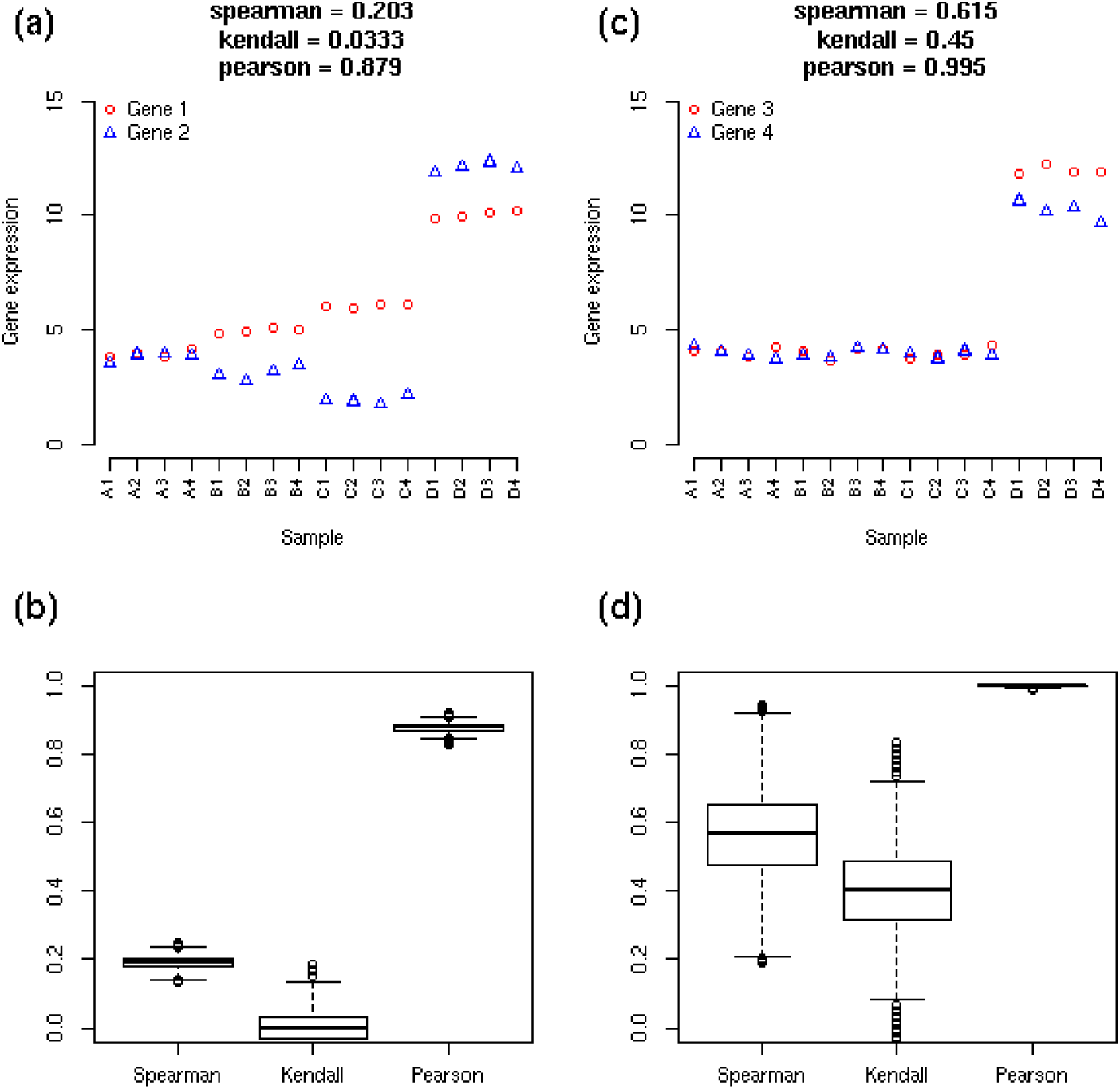
Simulated genes illustrating difficulties with correlation-based measures of co-expression.

Panel (b) shows the distribution of Spearman, Kendall and Pearson correlations over ten-thousand similarly simulated genes. Conversely, Panel (c) shows genes 3 & 4, which have similar expression over experimental conditions A, B and C and a similar change in expression in experimental condition D. However, panel (d) shows that the biological and technical variation between equivalently expressed samples in conditions A, B & C leads to highly unstable estimates of Kendall and Spearman correlation, implying that for many such genes co-expression will not be identifiable with these measures of correlation.

We thus propose a novel approach to generating clusters of co-expression from high throughput sequencing data by first considering clusters of ‘equivalent’ expression for a given gene across samples. If two genes share the same clustering of equivalent expression across samples, and the average expression between clusters is strictly monotonic between the two genes, then we can consider the genes to be co-expressed. For example, in Figure 1, genes 1 & 2 will share the same clustering (for the natural four cluster case), but the order of those clusterings by average gene expression is non-identical. Conversely, genes 3 & 4 also share the same clustering (clustering all A, B, & C samples together, with D forming a separate cluster) and the ordering of average gene expression is also preserved. We consequently seek to classify genes 3 & 4 as co-expressed and genes 1 & 2 as not co-expressed.

Based on this definition of co-expression, we develop two metrics defining the distance between genes from which clusters of co-expression can be produced. The first metric is based on an analysis of posterior likelihoods derived from empirical Bayesian analyses of RNA-seq data as implemented in the baySeq package (Hardcastle and Kelly, 2010). The second metric is based on comparisons between k-means clustering on the expression of individual genes.

## METHODS

We introduce two methods of generating clusters of co-expression from high-throughput sequencing data. These are both founded on the principle that two genes may be considered co-expressed if the observed values of the two events form identical equivalence clusters whose average behaviour is strictly monotonic between the two events.

### Likelihood derived distances

The first method follows on from a typical model-based analysis of RNA-seq data sets with the Bioconductor (Huber *et al.*, 2015) package baySeq (Hardcastle and Kelly, 2010). For relatively small (≥ 10) numbers of experimental conditions, we can compute posterior likelihoods of all possible combinations of differential behaviour between conditions. The distance metric between two genes *i* and *j* is then calculated as

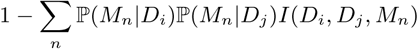

where ℙ(*M*_n_|*D*_*i*_) is the likelihood of model *M*_*n*_ given the observed data for gene *i* and *I*(*D*_*i*_, *D*_*j*_, *M*_*n*_) is an identity function which is 1 if the average expression of equivalently expressed (under model *M*_*n*_) samples for gene *i* is strictly monotonic with respect to the average expression of equivalently expressed (under model *M*_*n*_) samples for gene *j* and 0 otherwise. The product of the probabilities is then summed over all possible models.

### K-means derived distances

The second method developed depends on k-means clustering of the expression values across samples for each gene. For each gene, we assume an approximately log-normal distribution. Consequently, we first remove zeros by setting any zero read counts to one, and rescale the data by some library scaling factor to account for variation between library sizes (Bullard *et al.*, 2010) before transforming to the log_2_-scale.

For each gene, we use Tibshirani’s gap statistic () to determine whether any structure is present in the data or if expression across all samples may be considered equivalent. If expression is non-equivalent across samples, we cluster each gene using k-means clustering for *k* = 1 to *k* = *N*, where *N* is the number of samples. For each gene *i* and number of clusters *k*, we define the clustering as *C*_*ik*_ and calculate *d*_*ik*_, the maximum absolute difference between the expression of gene *i* in an individual sample and the centroid of the cluster to which that sample is assigned. To derive the clustering *C*_*ik*_ with minimum *d*_*ik*_, we repeat the k-means clusterings over one hundred iterations with randomly selected initial centroids. For each pair of genes *i* and *j*, the distance between them is calculated as

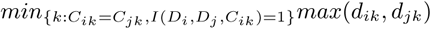

That is, over the values of *k* for which the clusterings *C*_*ik*_ and *C*_*ij*_ are identical, and for which the average expression of equivalently expressed (under the clustering *C*_*ik*_) samples for gene *i* is strictly monotonic with respect to the average expression of equivalently expressed (under the clustering *C*_*ik*_) samples for gene *j*, we choose the maximum of *d*_*ik*_ and *d*_*jk*_. This maximum represents the goodness of fit of the clustering *C*_*ik*_ for genes *i* and *j*. We then choose the minimum such distance over all possible values for *k*, noting that the conditions *C*_*ik*_ = *C*_*jk*_,*I*(*D*_*i*_, *D*_*j*_, *C*_*ik*_) = 1} will always hold for *k* = 1 and so this minimum is guaranteed to exist.

We note also that if *D*_*i*_ is strictly monotonic with respect to *D*_*j*_ (i.e., *D*_*i*_ and *D*_*j*_ are perfectly rank correlated) then *C*_*iN*_ = *C*_*jN*_, since each element of *D*_*i*_ and *D*_*j*_ will form a separate cluster, and *I*(*D*_*i*_, *D*_*j*_, *C*_*ik*_) = 1, with *d*_*ik*_ = *d*_*jk*_ = 0, and so the distance between genes *i* and *j* will be zero.

### Clustering

Given a metric defining distance between each pair of genes, clustering can be performed by any agglomerative method. If single-linkage agglomeration is used (the default proposed by the clusterSeq package) then for a given genei, only the nearest neighbouring gene *j* for *j* > *i* need be identified; consequently single-linkage agglomeration is relatively efficient when applied to large numbers of genes. However, it is also possible to construct the complete distance matrix for all genes and apply alternative methods of agglomeration.

Some threshold must be provided on the distances beyond which no further agglomeration occurs. We suggest as defaults a distance of 0.5 for the likelihood-derived metric; this threshold corresponds to the likelihood that two genes have less than a 50% likelihood of being equivalently expressed. We suggest a distance of 1 for the k-means-derived metric, which implies a 2-fold change or more between the greatest outlier of the best matched clusterings between two genes and the centroid of the cluster to which the outlier belongs.

## RESULTS

We test the clustering methods on a set of time series data in female rat thymus tissues (Yu *et al.*, 2014), in which samples were taken at 2, 6, 21 and 104 weeks. By identifying clusters of genes demonstrating similar patterns of expression over time we can identify time dependencies within the data.

Figure 2 shows the six largest clusters identified by clustering based on likelihood-derived distances. The largest cluster identified shows a loss of expression in 104 week-old rats, perhaps corresponding to the known phenomenon of thymic involution (Shanley *et al.*, 2009).

**Fig. 2.**
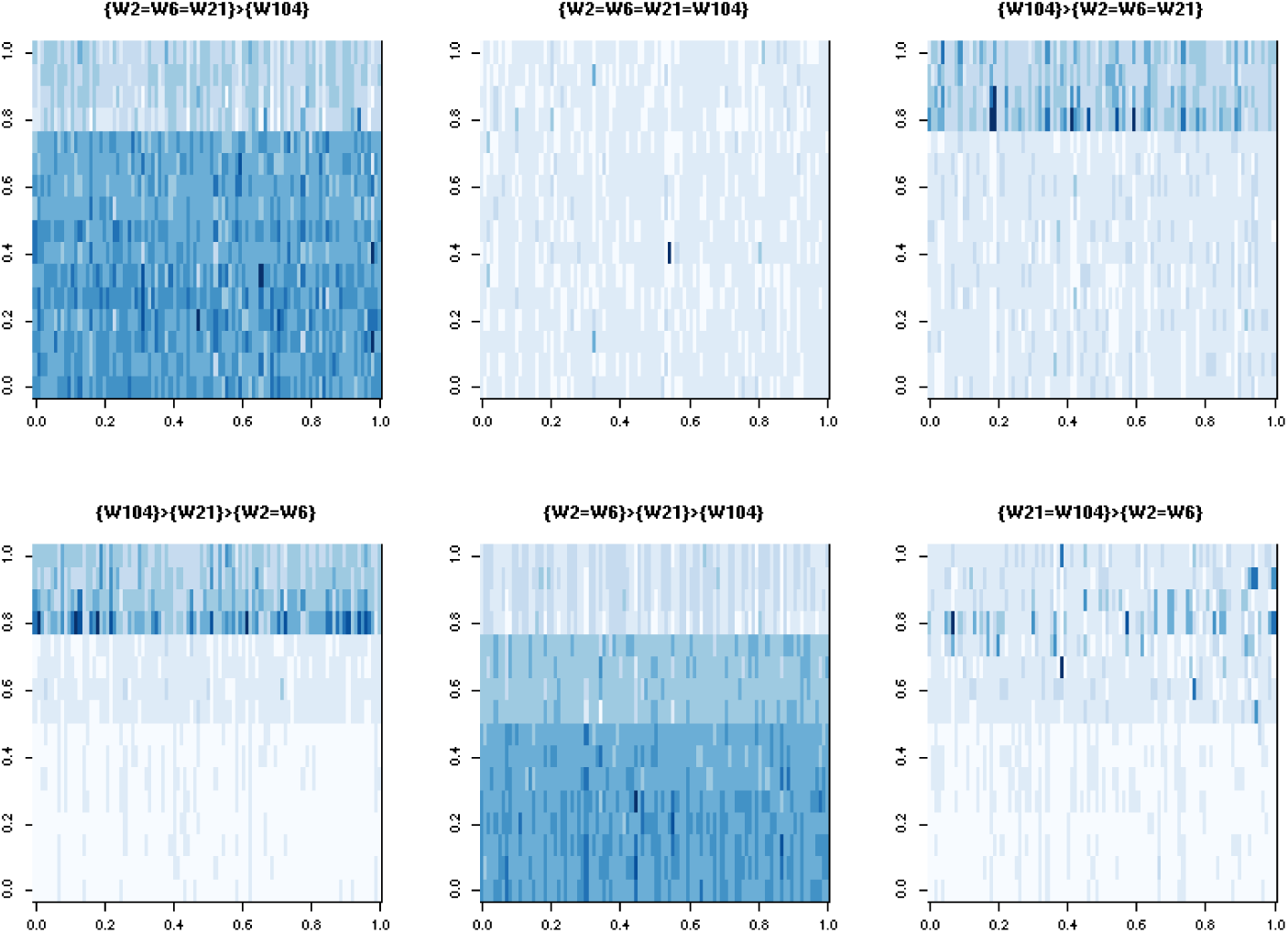
Heatmaps of the six largest clusters identified by likelihood-derived distance analyses.

## CONCLUSION

We have presented two novel methods for identifying clusters of co-expression that account for structures of equivalent expression within and between experimental conditions. These methods are released as the Bioconductor package clusterSeq (https://bioconductor.org/packages/release/bioc/html/clusterSeq.html)

The first method extends the empirical Bayesian analysis implemented in the baySeq package. This method is particularly well suited to experimental designs incorporating replication, as it is able to naturally incorporate the known replicate structure within the data. However, this method requires that the replicate structure is known in advance.

The second method uses a k-means approach to infer a clustering of samples for each gene before comparing these clusterings between genes to infer the distance between them. This method is applicable even without a known replicate structure, and scales well to detect co-expression in large numbers of samples. This approach has been used successfully to detect co-expression in the baseline experiments in the Expression Atlas (Petryszak *et al.*, 2014).

The methods developed here are demonstrated on gene expression data. However, it is relatively straightforward to generalise these analyses to identify co-expression in any high-throughput sequencing data. The generalisation of the empirical Bayesian methods described in Hardcastle (2016) allows posterior likelihoods of differential behaviour to be identified for any quantifiable biomolecular event. Similarly, the k-means clustering of gene expression data can be adapted to, for example, cluster proportions of methylation derived from methylation loci.

## ACKNOWLEDGEMENTS

TJH is supported by European Research Council Advanced Investigator Grant ERC-2013-AdG 340642 - TRIBE. IP has been funded by the National Science Foundation of USA grant to Gramene database [NSF IOS #1127112] and by the European Molecular Biology Laboratory (EMBL) member states.

